## Supplementary material for "Cost-benefit Tradeoff Mediates the Rule- to Memory-based Processing Transition during Practice"

**Supplementary Notes**

***Note S1.*** *Behavioral Analysis with All Cued Positions.*

We observed that cued position 5 showed better performance compared to cued position 4 and that cued position 1 outperformed all other cued positions. These are possibly due to better memory of the first and last items of a sequence [1]. To test the robustness of the behavioral findings, we repeated the mixed-effect model analyses in the main text while including all five cued positions. That is, we used cued position (1–5), block (1–6) and their interaction as predictors of performance. We found similar results as reported in the main text. Specifically, the RT results showed a significant main effect of block ($\beta$ = –0.36, SE = 0.03, 95% confidence interval (CI) = [–0.42 –0.31], $t(33.0) = -13.80$, $p < .001$, $\eta_p^2 = .27$), a significant main effect of position ($\beta$ = 0.29, SE = 0.04, 95% CI = [0.21 0.37], $t(33.0) = 7.41$, $p < .001$, $\eta_p^2 = .19$), and a significant interaction effect ($\beta$ = –0.09, SE = 0.02, 95% CI = [–0.13 –0.05], $t(580.8) = -4.47$, $p < .001$, $\eta_p^2 = .02$). Post-hoc comparisons showed that the slope of position in block 1 was larger than block 3, 4, 5, and 6, whereas block 2 was larger than block 6, *p*s < .05. The ER results showed a significant main effect of block ($\beta$ = –0.23, SE = 0.05, 95% CI = [–0.33 –0.13], $t(33.0) = -4.78$, $p < .001$, $\eta_p^2 = .07$), position ($\beta$ = 0.16, SE = 0.04, 95% CI = [0.08 0.24], $t(38.8) = 4.09$, $p < .001$, $\eta_p^2 = .04$), and a significant interaction effect ($\beta$ = –0.13, SE = 0.04, 95% CI = [–0.21 –0.05], $t(39.6) = -3.45$, $p < .001$, $\eta_p^2 = .02$). Post-hoc comparisons suggested that the slope of position in block 1 was larger than block 3, 4, and 6, whereas block 2 was larger than block 3 and 6, and block 4 and block 5 were larger than block 6, *p*s < .05. These results suggest that our behavioral findings were not qualitatively affected by the exclusion of cued position 5.

***Note S2.*** *No Evidence Supporting the Discount of Rule-based Strategy Cost in the Cost-benefit Analysis*

With a new model (M6), we tested the possibility that practicing the rule-based strategy would reduce its cost in the cost-benefit analysis. To this end, the rule-based strategy cost was multiplied by a discount factor (D), which takes the following form:

$$D = e^{\delta N},$$

where N denotes the number of experienced rule trials (including those occurring during the transition test in the training phase). $\delta$ is a free parameter that ranges from –1000 to 0, which simulates a wide range of decrease trajectories, including non-linear decrease and quasi-linear decrease with various slopes.

M6 was constructed by adding the discount factor to our previously validated model (M0). Model fitting resulted in a group sum Akaike information criterion (AIC) of –3928, higher than that of M0 (–4019), $p < .001$. This result suggests that the acceleration of sequence recall with practice does not significantly affect the cost-benefit analysis.

***Note S3.*** *No Evidence Supporting Backward Replay*

We investigated whether participants engaged in backward replay process [2] (i.e., starting by recalling the 5$^{th}$ position and then transitioning backward until the cued position). One might intuitively expect this to occur for later positions such as A4, since executing a backward replay only requires one step, whereas a forward replay demands three steps. However, given that our training exclusively focused on forward transitions, it remains unclear whether participants would spontaneously adopt backward replay strategies during testing. To address this question, we developed a new model (M7). With this model, we included in the cost-benefit analysis the value of implementing backward replay for each trial. To capture the individual differences in the cost of backward replay, the cost per replaying step of backward replay is that of the forward replay scaled by a factor of $\frac{1}{\mu}$, where $\mu$ ranges from 0 to 1. In other words, for a single step of replay, the cost of backward replay ranges from the cost of forward replay (when $\mu = 1$) to infinite (when $\mu = 0$). We opted for this implementation under the assumption that backward replay is more costly than or as costly as forward replay. Our results showed that this model performed worse (AIC = –3984) compared to M0 (AIC = –4019), $p < .001$. Further examination of M7 revealed that, on average, participants were estimated to engage in backward replay for 5.9 trials, whereas forward replay was implemented for 80.8 trials, suggesting that, even if backward replay is employed, it is far less prevalent among participants than forward replay.

***Note S4.*** *Detailed Task Design in the Training Phase.*

The training task was programmed with JsPsych (version 7.0, https://www.jspsych.org/) and conducted online. The training task aimed to facilitate participants in learning the transitions within two task sequences. At the beginning of each mini-block, participants were instructed to equip an avatar based on a goal image (Fig. 1A) in five steps (trials). We used two sequences, coded A and B, with the order of a→b→c→d→e and d→c→a→e→b, respectively. Each lowercase letter represents an equipment type (HELMET, VEST, GLASSES, WEAPON, or TOOL). The letter-equipment type mapping was randomly chosen for each participant. For each mini-block, the goal image was a fully equipped avatar with all gears randomly selected, so that the participants would learn the order of equipment type rather than specific gears.

Each trial started with the presentation of a sequence cue (A or B) for 2s, followed by a fixation cross for 2s and the presentation of the goal image for 1s. After a blank screen of 0.5s, a partially equipped avatar (representing current progress) and two gears of the same type were presented. Participants were required to choose the gear from the goal image using a button press (left or right) within 2.5s, followed by feedback. The same trial was repeated if an error was committed. After the fifth trial, a “goal achieved” screen was shown for 2s to signal the end of the mini-block.

Following each mini-block, we tested the participant's memory of task transitions with two test trials (Fig. S1). On each test trial, participants needed to choose the 1$^{st}$/2$^{nd}$/3$^{rd}$ task following a cued task. The cued task was selected randomly from the first four tasks from each sequence. The number of transitions (1, 2, or 3) was randomly sampled from a geometric distribution (p = 0.5). Participants responded using keys "Q", "W", "O" and "P" with no time limit. Incorrect responses were followed by feedback and repetition of the test trial. The average accuracy for the test trials was displayed at the end of each block.

Participants performed 3-6 blocks of 12 mini-blocks each until their block-average accuracy of test trials reached 90% in the last block. Participants who could not meet the 90% accuracy after six blocks were not scanned. Thirty-five participants underwent the training phase 4-24 hours before scanning. They repeated the training phase before scanning until they could reach the 90% accuracy threshold again (all completed within 2 blocks). One participant completed the training phase immediately before scanning.

***Note S5.*** *fMRI Data Preprocessing*

Results included in this manuscript come from preprocessing performed using fMRIPrep 22.0.2 (RRID:SCR_016216) [3,4], which is based on Nipype 1.8.5 (RRID:SCR_002502) [5,6].

*Preprocessing of B0 inhomogeneity mappings.* A B0-nonuniformity map (or fieldmap) was estimated based on two (or more) echo-planar imaging (EPI) references with topup (FSL 6.0.5.1:57b01774) [7].

*Anatomical data preprocessing.* The T1-weighted (T1w) image was corrected for intensity non-uniformity (INU) with N4BiasFieldCorrection [8], distributed with ANTs 2.3.3 (RRID:SCR_004757) [9], and used as T1w-reference throughout the workflow. The T1w-reference was then skull-stripped with a Nipype implementation of the antsBrainExtraction.sh workflow (from ANTs), using OASIS30ANTs as target template. Brain tissue segmentation of cerebrospinal fluid (CSF), white-matter (WM) and gray-matter (GM) was performed on the brain-extracted T1w using fast (FSL 6.0.5.1:57b01774, RRID:SCR_002823) [10]. Volume-based spatial normalization to one standard space (MNI152NLin2009cAsym) was performed through nonlinear registration with antsRegistration (ANTs 2.3.3), using brain-extracted versions of both T1w reference and the T1w template. The following template was selected for spatial normalization: ICBM 152 Nonlinear Asymmetrical template version 2009c [RRID:SCR_008796; TemplateFlow ID: MNI152NLin2009cAsym] [11].

*Functional data preprocessing.* For each of the 6 BOLD runs, the following preprocessing was performed. First, a reference volume and its skull-stripped version were generated using a custom methodology of fMRIPrep. Head-motion parameters with respect to the BOLD reference (transformation matrices, and six corresponding rotation and translation parameters) are estimated before any spatiotemporal filtering using mcflirt (FSL 6.0.5.1:57b01774) [12]. The estimated fieldmap was then aligned with rigid-registration to the target EPI (echo-planar imaging) reference run. The field coefficients were mapped on to the reference EPI using the transform. The BOLD reference was then co-registered to the T1w reference using mri_coreg (FreeSurfer) followed by flirt (FSL 6.0.5.1:57b01774) [13] with the boundary-based registration [14] cost-function. Co-registration was configured with six degrees of freedom. Several confounding time-series were calculated based on the preprocessed BOLD: framewise displacement (FD), DVARS and three region-wise global signals. FD was computed using two formulations following Power (absolute sum of relative motions) [15] and Jenkinson (relative root mean square displacement between affines) [12]. FD and DVARS are calculated for each functional run, both using their implementations in Nipype (following the definitions by [15]). The three global signals are extracted within the CSF, the WM, and the whole-brain masks. Additionally, a set of physiological regressors were extracted to allow for component-based noise correction (CompCor) [16]. Principal components are estimated after high-pass filtering the preprocessed BOLD time-series (using a discrete cosine filter with 128s cut-off) for the two CompCor variants: temporal (tCompCor) and anatomical (aCompCor). tCompCor components are then calculated from the top 2% variable voxels within the brain mask. For aCompCor, three probabilistic masks (CSF, WM and combined CSF+WM) are generated in anatomical space. The implementation differs from that of Behzadi et al. in that instead of eroding the masks by 2 pixels on BOLD space, a mask of pixels that likely contain a volume fraction of GM is subtracted from the aCompCor masks. This mask is obtained by thresholding the corresponding partial volume map at 0.05, and it ensures components are not extracted from voxels containing a minimal fraction of GM. Finally, these masks are resampled into BOLD space and binarized by thresholding at 0.99 (as in the original implementation). Components are also calculated separately within the WM and CSF masks. For each CompCor decomposition, the k components with the largest singular values are retained, such that the retained components' time series are sufficient to explain 50 percent of variance across the nuisance mask (CSF, WM, combined, or temporal). The remaining components are dropped from consideration. The head-motion estimates calculated in the correction step were also placed within the corresponding confounds file. The confound time series derived from head motion estimates and global signals were expanded with the inclusion of temporal derivatives and quadratic terms for each [17]. Frames that exceeded a threshold of 0.5 mm FD or 1.5 standardized DVARS were annotated as motion outliers. Additional nuisance timeseries are calculated by means of principal components analysis of the signal found within a thin band (crown) of voxels around the edge of the brain, as proposed by ref [18]. The BOLD time-series were resampled into standard space, generating a preprocessed BOLD run in MNI152NLin2009cAsym space. First, a reference volume and its skull-stripped version were generated using a custom methodology of fMRIPrep. All resamplings can be performed with a single interpolation step by composing all the pertinent transformations (i.e., head-motion transform matrices, susceptibility distortion correction when available, and co-registrations to anatomical and output spaces). Gridded (volumetric) resamplings were performed using antsApplyTransforms (ANTs), configured with Lanczos interpolation to minimize the smoothing effects of other kernels [19]. Non-gridded (surface) resamplings were performed using mri_vol2surf (FreeSurfer).

Many internal operations of fMRIPrep use Nilearn 0.9.1 (RRID:SCR_001362) [20], mostly within the functional processing workflow. For more details of the pipeline, see the section corresponding to workflows in fMRIPrep's documentation.

*Copyright Waiver.* The above boilerplate text was automatically generated by fMRIPrep with the express intention that users should copy and paste this text into their manuscripts *unchanged*. It is released under the CC0 license.

***Note S6.*** *Univariate Evidence for Rule- and Memory-based Strategies.*

To detect the brain regions whose fMRI activation reflects key components of the two strategies, we conducted three LME models with ROI-averaged fMRI activation levels. The first LME model was conducted on rule trials and incorporated several fixed effect regressors, including the rule implementation cost C, $C*Times_R$ (to account for the acceleration of rule-based strategy due to practice), and the logarithm of trial number (to account for strategy-agnostic practice effect). The second LME model was conducted on memory trials and included fixed effect regressors of memory strength and the logarithm of trial number. The third LME model was conducted on both rule and memory trials and included fixed effect regressors of the decision variable (defined by the utility difference of alternative strategies, log-transformed), C, $C*Times_R$, memory strength, and the logarithm of trial number. The first two models also included an intercept, and the third model included two intercepts indicating the two trial types. Additionally, the intercepts and the slope of each regressor were also treated as random effects at the subject level. All predictors were standardized. Statistical results were corrected for multiple comparisons across all the ROIs with false discovery rate (FDR) at the level of $q < 0.05$, with each individual ROI at uncorrected $p < .001$, both one-tailed for the rule trials and two-tailed for the memory trials based on our different hypotheses (see below).

We predicted stronger fMRI activation when later positions were cued on rule trials, possibly due to increased effort in implementing the rule-based strategy (e.g., the steps of replay [21], difficulty, and working memory load). Higher rule implementation costs were accompanied by stronger activation in several frontal regions, including Human Connectome Project (HCP) Atlas [22] areas 8C, anterior inferior frontal junction (IFJa), posterior 47 rostral (p47r), anterior 47 rostral (a47r), anterior 9-46 ventral (a9-46v), posterior 32 prime (p32pr), inferior 6-8 (i6-8) in the left hemisphere and 8C, posterior 9-46 ventral (p9-46v), 44 in the right hemisphere, *p*s < 0.05 corrected (Fig. S3A-B). On memory trials, fMRI activation may be stronger when the cue-task association is stronger (reflecting the memory retrieval effect) [23] or weaker (reflecting the uncertainty effect) [24]. We found stronger brain activation with stronger memory of the cued position-task association in the left PF (Fig. S3C-D, $p < 0.05$ corrected), and stronger brain activation with weaker memory of the cued position-task association in the left supplementary and cingulate eye field (SCEF) and the left posterior IFJ (IFJp) (Fig. S3E-F). No other regions survived the correction for multiple comparisons. However, the decision variable was not associated with any specific brain regions, suggesting that the encoding of the cost-benefit tradeoff in the dorsomedial prefrontal cortex might only occur in a distributed fashion (Fig. 5C). These results are consistent with previous findings [24-26] and provide a sanity check of the fMRI data.

***Note S7.*** *Detailed RSA Methods.*

*Analysis 1: Strategy-specific Representations on Rule and Memory Trials.*

The computational model categorized trials into rule and memory trials based on behavioral data. We then tested if this categorization accounted for the fMRI data. If the cost-benefit analysis decides which strategy to implement, we would expect a stronger rule effect on the rule than the memory trials and a stronger memory effect on memory than rule trials. To this end, we conducted RSA including the two factors (rule/memory effects as illustrated in Fig. 3 and rule/memory trials classified by the computational model). The RSA measures the fMRI activation pattern similarity between two trials. This measure serves as the dependent variable. The key independent variables include the rule and memory effects encoding hypothetical similarity based on the corresponding strategy. Specifically, the rule effect is operationalized as the number of shared tasks between two trials (counting from the beginning of the sequence) excluding the cued task (Fig. 3B). The memory effect encodes whether the two trials share the same cue, which captures the memory strategy (i.e., directly retrieving the answer from the cue without engaging the rule-based strategy, Fig. 3C). Each effect was coded separately within each trial type to test the difference in the effect between rule and memory trials. To account for the potential context effect (i.e., a higher similarity for within-sequence trial pairs than across-sequence trial pairs), trial pairs were further divided into within-sequence and across-sequence trial pairs. We further added separate intercepts to each trial type. As a result, the model is formulated as follows:

$$RSM_{brain} = \beta_0 R1 + \beta_1 R2 + \beta_2 M1 + \beta_3 M2 + \beta_4 RSM_{Rule\_R} + \beta_5 RSM_{Rule\_M} + \beta_6 RSM_{Cue\_R1} + \beta_7 RSM_{Cue\_R2} + \beta_8 RSM_{Cue\_M1} + \beta_9 RSM_{Cue\_M2} + \beta_{10} RSM_{Univoxel\_trial1} + \beta_{11} RSM_{Univoxel\_trial2} + (R1|subject) + (R2|subject) + (M1|subject) + (M2|subject) + RandomSlopes + \varepsilon.$$

In this equation, $RSM_{brain}$ represents the representational similarity matrix (RSM) derived from fMRI activation patterns. M and R encode memory and rule trials, respectively. Rule and Cue in subscript encode their corresponding effects. The division of trial pairs based on the context is reflected by the R1 (rule trial pairs from the same sequence), R2 (rule trial pairs from different sequences), M1 (memory trial pairs from the same sequence), and M2 (memory trial pairs from different sequences), respectively. These regressors were added to account for potential differences in the pattern similarity between same- and different-sequence trial pairs and between different trial types. $RSM_{Rule}$ and $RSM_{Cue}$ encode hypothesized pattern similarity levels based on rule- and memory-based strategies, respectively (Fig. 3B, C). The term RandomSlopes represents the collection of random effects for the slope of each fixed effect. Additionally, the $RSM_{Univoxel\_trial1}$ and $RSM_{Univoxel\_trial2}$ capture the univoxel activation for each trial of the pair. All random effects were set at the subject level. An illustration of this model can be found in Fig. S5.

To further test the difference in rule and memory effects between the rule trials and memory trials from the same sequence, we conducted three contrast analyses. These included (1) testing if the rule effect was stronger in rule trials compared to memory trials ($\beta_4 > \beta_5$), (2) testing if the cue effect from the same sequence was stronger in memory trials compared to rule trials ($\beta_8 > \beta_6$), and (3) testing if there were regions showing a double dissociation between rule/memory strategy effect and trial type using a conjunction analysis of (1) and (2).

*Analysis 2: fMRI Boundary Effects around Transition Points*

This analysis aimed to assess changes in activation patterns during strategy transition. As in the behavioral and univariate analyses, here we focused on the two trials immediately adjacent to a transition point. We denoted the transition point at t and coded the last rule trial as t–1 and the first memory trial as t+1. We computed the similarity between these two trials and each of the trials 2–6 steps before/after the transition point (baseline trials). We hypothesized a decrease in pattern similarity when trials were from opposite sides of the boundary (i.e., transition point), such that trial t–1 would show greater similar to the rule baseline trials (within trial type) compared to the memory baseline trials (between trial types), and vice versa for trial t+1. Thus, we constructed a regressor representing whether two trials are of the same trial type ($RSM_{Trialtype}$). To account for the potential confound of temporal proximity, we included a nuisance regressor encoding temporal distance between each trial pair ($RSM_{Distance}$). As above, trial-wise univariate activation levels were included as additional nuisance regressors. The resulting LME model takes the following form:

$$RSM_{brain} = \beta_0 + \beta_1 RSM_{Trialtype} + \beta_2 RSM_{Distance} + \beta_3 RSM_{Univoxel\_trial1} + \beta_4 RSM_{Univoxel\_trial2} + RandomSlopes + \varepsilon.$$

All trial pairs in this analysis were limited to different runs to control for RSA confounds from within-run trial pairs while limiting the analysis to trials close to transition points [27].

*Analysis 3: Pattern Separation.*

The use of the memory strategy predicted that the learning of cue-task associations would prompt distinctive patterns to emerge, effectively separating the conditions [28]. To test this prediction of increased pattern separation with practice, we computed the representational similarity between trial pairs that were the same temporal distance away from their respective transition points. To better capture the temporal trajectory of pattern separation for each cue, the temporal distance between a trial and its transition point counts only trials with the same cue. To test potential changes in the degree of pattern separation before and after strategy transition, we introduced two separate regressors in our model: one for rule trials and another for memory trials. As above, we also added the nuisance regressors of trial distance and trial-wise univariate activations to the LME model:

$$RSM_{brain} = \beta_0 M + \beta_1 R + \beta_2 RSM_M + \beta_3 RSM_R + \beta_4 RSM_{Distance} + \beta_5 RSM_{Univoxel\_trial1} + \beta_6 RSM_{Univoxel\_trial2} + RandomSlopes + \varepsilon.$$

This analysis focused on the average effect of trial number across both rule ($\beta_2$) and memory ($\beta_3$) trials. This allowed us to assess the level of pattern separation in relation to the temporal distance to the transition points. We also tested $\beta_2$- $\beta_3$ to explore whether the speed of pattern separation changed following strategy transition.

***Note S8.*** *Evidence of Replay in Rule-based Task Retrieval.*

We hypothesized that rule-based processing involves replaying tasks within the same sequence to facilitate the retrieval of the cued task. To test this, we examined whether tasks preceding the cued position could be decoded above chance level through a decoding analysis. For each task, we trained a lasso logistic regression classifier to predict whether the task was replayed on each trial (i.e., from the beginning of the sequence to the cued task), setting the regularization coefficient at 0.006 [29]. To mitigate overfitting, the leave-one-block-out cross-validation was applied. The trained classifiers were then tested on the left-out data to obtain a probabilistic prediction of the presence of each task on each trial. As a result, given the fMRI activation pattern of a trial, the classifier produced a probabilistic prediction for each task being replayed. We used these predictions to perform two further regression analyses:

The first analysis aimed to examine if the replayed tasks could be decoded (i.e., showing a higher predicted probability of being replayed than chance level) from rule trials but not memory trials. We conducted two separate analyses for the two trial types using the same formula but different dependent variables:

$$Probability\_prediction\ (R\ or\ M) = \beta_0 + \beta_1 Replayed\_tasks + \beta_2 Cued\_task + \beta_3 Other\_task + RandomSlopes + \varepsilon,$$

where Probability_prediction is a trial × task matrix containing probability values, Replayed_tasks is a binary predictor indicating the hypothesized replayed tasks (i.e., tasks prior to the cued position) on each trial, Cued_task denotes the cued task, and Other_task is another binary predictor denoting the tasks posterior to the cued position.

To avoid potential bias in the decoding process, we established a baseline decoding performance by conducting permutation tests. Permutations were done by randomly shuffling the labels (e.g., A4) of each trial 500 times. Results showed that, for the 63 regions showing rule implementation representations (Fig. 4A), 4 regions (left STSd_posterior, left TE1_anterior, left IFSa, and right TE2_posterior) showed a significant Replayed_tasks effect on rule trials ($ps < 0.05$, FDR corrected), suggesting that the hypothesized replayed tasks can be decoded above chance level. In comparison, no regions showed a Replayed_tasks effect on memory trials, $ps = 1$, uncorrected.

To further test whether the Replayed_tasks effect was stronger on rule trials than on memory trials, we conducted an interaction analysis with the following formula:

$$\text{Probability\_prediction} = \beta_0 + \beta_1\text{Replayed\_tasks} * \text{Trialtype} + \beta_2\text{Cued\_task} * \text{Trialtype} + \text{RandomSlopes} + \varepsilon,$$

where Trialtype is a categorical regressor denoting the trial type (1 for rule-based trials and –1 for memory-based trials). Results showed that, among the 108 regions identified in Fig. 4C, eighty-one regions showed this effect, *p*s < 0.05, FDR corrected (Fig. S7A). Across the 426 whole-brain regions, we also identified similar brain distributions with FDR correction (Fig. S7B).

### Supplementary Figures

Fig. S1

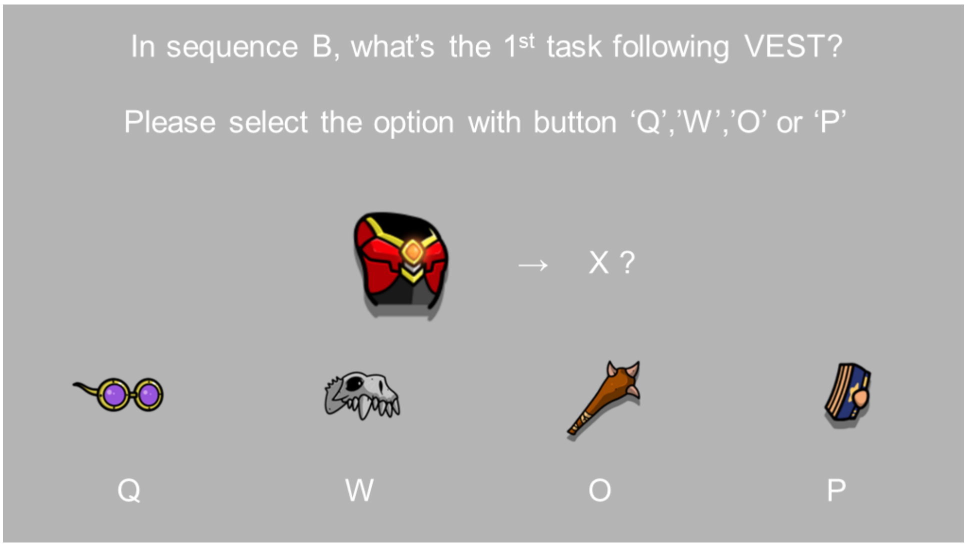


**Fig. S1. Design of the test task during the training phase.** This task aimed to test participants' memory of task transitions. They were asked to choose the $1^{st}$/$2^{nd}$/$3^{rd}$ tasks (in this example, the $1^{st}$ task) following a prompted task within one sequence (in this example, sequence "B"). Responses were made with "Q", "W", "O" or "P" buttons on the keyboard without time constraints.

Fig. S2

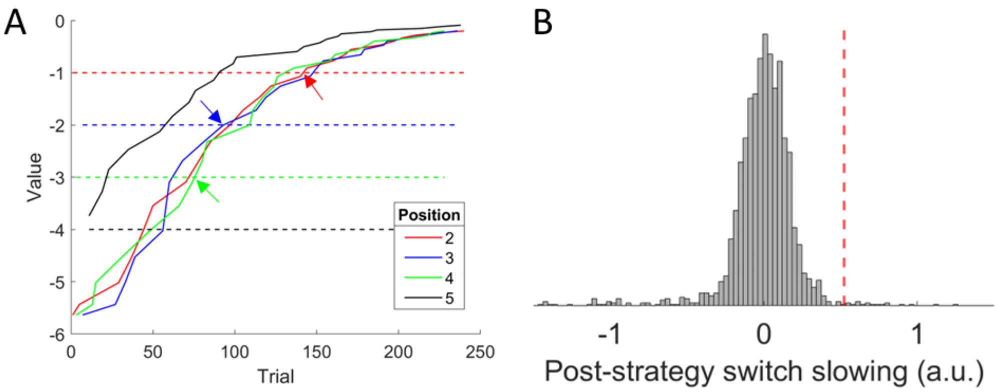


**Fig. S2. Model estimation of strategy transition.** A) Strategy transition based on the value (i.e., cost-benefit tradeoff) between the rule-based strategy (dashed lines) and the memory strategy (solid lines) in an exemplar sequence of a subject. The arrows indicate the transition points for different cued positions. The value for the rule-based strategy remains constant over time, thus the dashed lines are horizontal. On the other hand, cue-task associations strengthen over learning, thus the solid line increases monotonically. A strategy switch occurs when the two lines cross (i.e., when the value of the memory strategy becomes better than that of the rule-based strategy). Note that position 5 has a higher value for memory than for the rule-based strategies throughout all trials, so the participant applied the memory strategy from the beginning. B) The permutation results (histogram) for the post-strategy switch slowing obtained by randomizing the model parameters. This analysis seeks to examine whether the post-transition slowing is specific to the transition points identified by the model. The red dashed line shows the coefficient derived from real data and is significantly higher than the null distribution, $p$ = .005.

Fig. S3

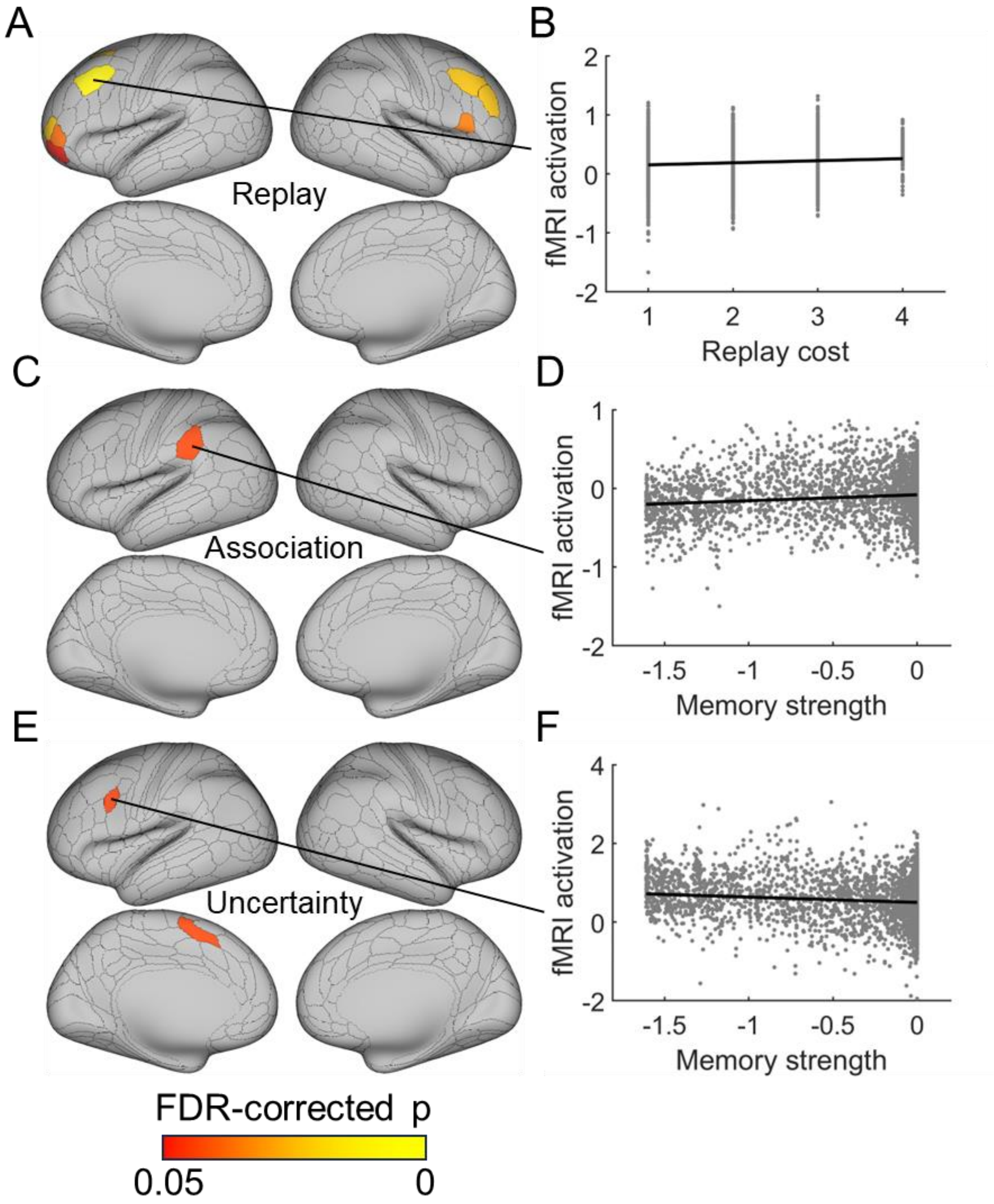


**Fig. S3. Frontoparietal fMRI activation for rule- and memory-based strategies.** A) Regions showing higher fMRI activation with higher rule implementation cost (i.e., number of steps replayed). B) Trial-wise fMRI activation in the left 8C region as a function of rule implementation cost. C) and E) show regions with higher trial-wise activation for stronger memory strength (i.e., cue-task associations) and higher trial-wise activation for lower memory strength (i.e., higher uncertainty), respectively. D) and F) show the trial-wise fMRI activation in the left PF and left IFJp, respectively, plotted as a function of memory strength quantified as reversed entropy.

Fig. S4

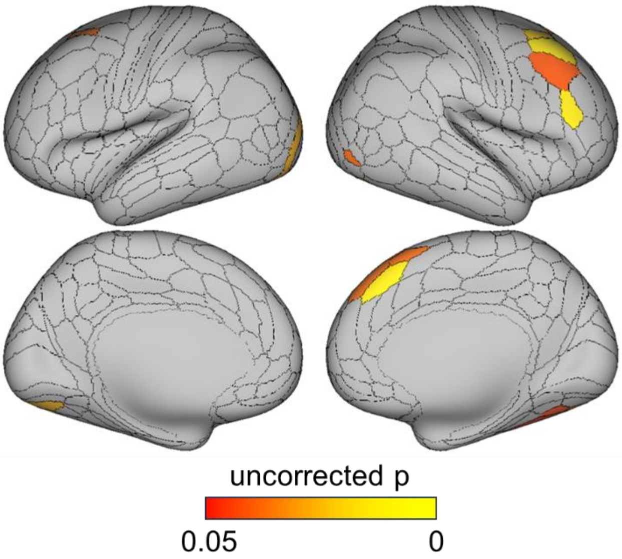


**Fig. S4. Univariate strategy transition cost.** Displayed are regions showing stronger fMRI activation on the first memory trial compared to the last rule trial, thresholded at uncorrected $p < .05$. The involvement of frontal regions is consistent with the switching cost hypothesis that occurs when transferring from one task to another [30,31].

Fig. S5

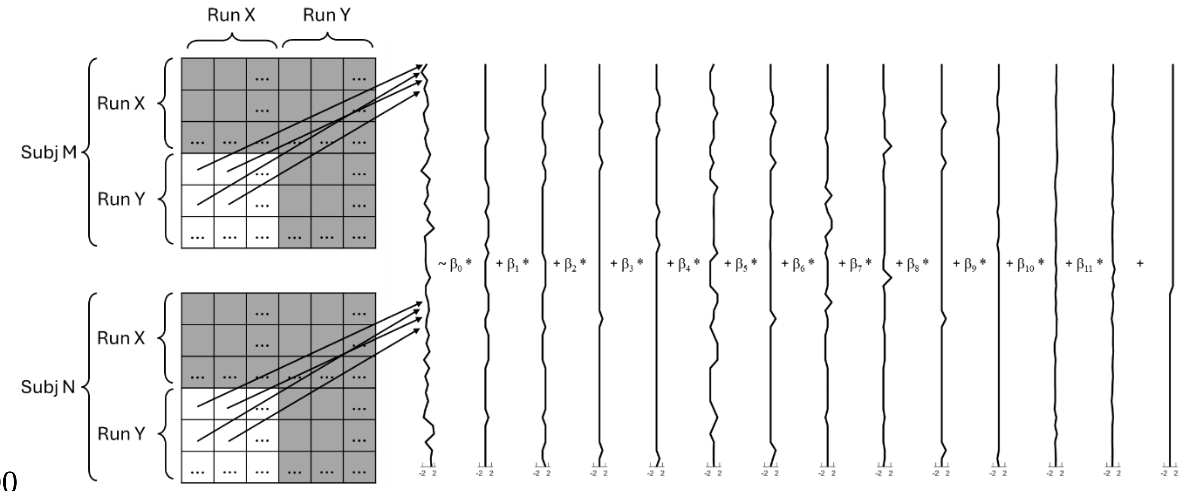


Fig. S5. **LME structure demonstration for RSA Analysis 1 (detailed in Note S6).** Sample data from two participants is plotted to illustrate how the values within the representational similarity matrices are used in the LME. In the left panel, the white cells, representing the similarities/differences in a model variable between two trials, are converted into a vector in the LME, while the gray-colored cells are removed due to either within-run correlation (diagonal) or duplication (upright off-diagonal). In the right panel, the first column represents z-transformed representational similarity values. Columns 2 through 13 correspond to regressors for different experimental conditions: R1, R2, M1, M2, $RSM_{Rule\_R}$, $RSM_{Rule\_M}$, $RSM_{Cue\_R1}$, $RSM_{Cue\_R2}$, $RSM_{Cue\_M1}$, $RSM_{Cue\_M2}$, $RSM_{Univoxel\_trial1}$ and $RSM_{Univoxel\_trial2}$. The participant identifier is represented in the final column.

Fig. S6

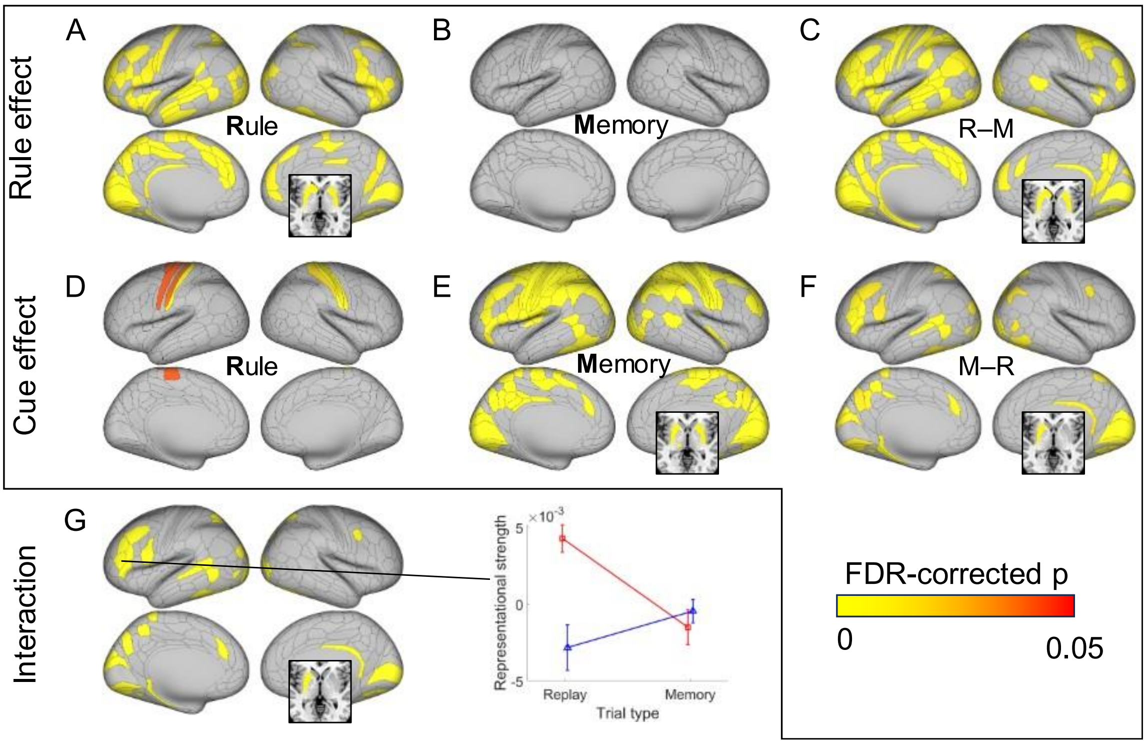


Fig. S6. **Double dissociation results with models controlling for difficulty.** A significant representation of the rule effect is observed on rule (R) trials (panel A) but not on memory (M) trials (panel B). C) A stronger rule effect on rule than memory trials (R–M) was found in frontoparietal, temporal, occipital, and subcortical regions. In contrast, the cue effect is exhibited in more regions on memory trials (panel E) than on rule trials (panel D), and regions showing a stronger memory effect on memory than rule trials (M–R) include frontoparietal, temporal, occipital, and subcortical regions (panel F). G) Regions showing double dissociation (i.e., the conjunction of C and F) with both stronger rule effect representation on rule trials and stronger cue effect representation on memory trials. All subcortical regions are depicted in an axial slice at z = 0. The right panel illustrates the double dissociation with an example region (left IFSa).

Fig. S7

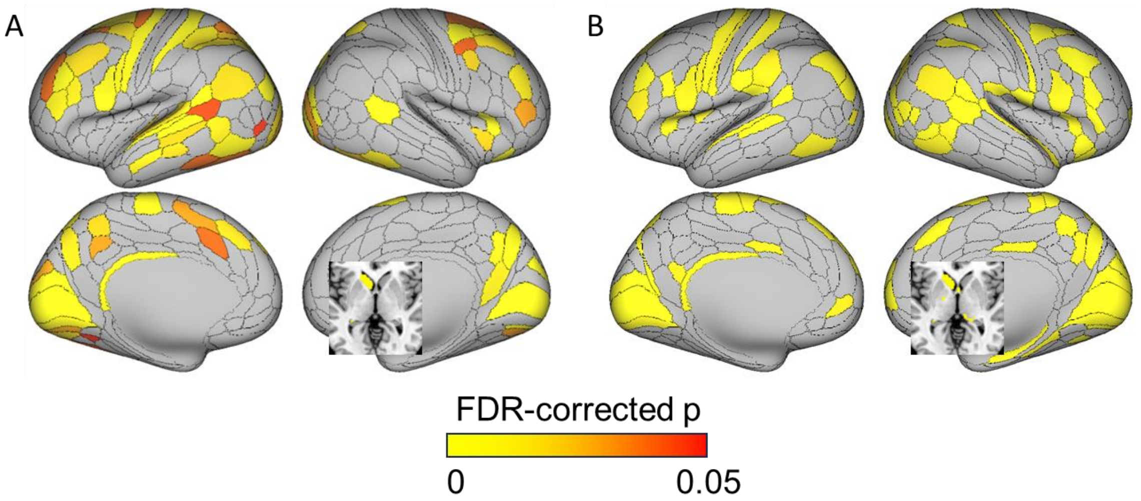


Fig. S7. **Decoding evidence supporting task replay.** Both panels show regions identified with stronger decoding of replayed tasks on rule than memory trials. A) FDR-corrected results using regions identified in Fig. 4C as search regions. B) FDR-corrected results using whole-brain ROIs.

### Supplementary Tables

***Table S1.*** *A detailed list of regions showing rule effects, cue effects, and the double dissociation interaction effects that are displayed in Figure 5 (see the Excel file "Table S1").*

### Supplementary References


1. Murdock, B.B. (1962). The serial position effect of free recall. Journal of Experimental Psychology *64*, 482-488. 10.1037/h0045106.
2. Foster, D.J., and Wilson, M.A. (2006). Reverse replay of behavioural sequences in hippocampal place cells during the awake state. Nature *440*, 680-683. 10.1038/nature04587.
3. Esteban, O., Markiewicz, C.J., Blair, R.W., Moodie, C.A., Isik, A.I., Erramuzpe, A., Kent, J.D., Goncalves, M., DuPre, E., Snyder, M., et al. (2019). fMRIPrep: a robust preprocessing pipeline for functional MRI. Nat Methods *16*, 111-+. 10.1038/s41592-018-0235-4.
4. Esteban, Oscar, Blair, R., Markiewicz, C.J., Berleant, S.L., Moodie, C., Ma, F., and Ayse Ilkay Isik, e.a. (2018). FMRIPrep (Software).
5. Gorgolewski, K., Burns, C.D., Madison, C., Clark, D., Halchenko, Y.O., Waskom, M.L., and Ghosh, S.S. (2011). Nipype: a flexible, lightweight and extensible neuroimaging data processing framework in python. Frontiers in neuroinformatics *5*, 13. 10.3389/fninf.2011.00013.
6. Gorgolewski, J., K., Esteban, O., Markiewicz, C.J., Ziegler, E., Ellis, D.G., Notter, M.P., and al., D.J.e. (2018). Nipype (Software).
7. Andersson, J.L., Skare, S., and Ashburner, J. (2003). How to correct susceptibility distortions in spin-echo echo-planar images: application to diffusion tensor imaging. Neuroimage *20*, 870-888. 10.1016/S1053-8119(03)00336-7.
8. Tustison, N.J., Avants, B.B., Cook, P.A., Zheng, Y., Egan, A., Yushkevich, P.A., and Gee, J.C. (2010). N4ITK: improved N3 bias correction. IEEE Trans Med Imaging *29*, 1310-1320. 10.1109/TMI.2010.2046908.
9. Avants, B.B., Epstein, C.L., Grossman, M., and Gee, J.C. (2008). Symmetric diffeomorphic image registration with cross-correlation: evaluating automated labeling of elderly and neurodegenerative brain. Med Image Anal *12*, 26-41. 10.1016/j.media.2007.06.004.
10. Zhang, Y., Brady, M., and Smith, S. (2001). Segmentation of brain MR images through a hidden Markov random field model and the expectation-maximization algorithm. IEEE Trans Med Imaging *20*, 45-57. 10.1109/42.906424.
11. Fonov, V.S., Evans, A.C., McKinstry, R.C., Almli, C.R., and Collins, D.L. (2009). Unbiased nonlinear average age-appropriate brain templates from birth to adulthood. NeuroImage *47*. 10.1016/s1053-8119(09)70884-5.
12. Jenkinson, M., Bannister, P., Brady, M., and Smith, S. (2002). Improved optimization for the robust and accurate linear registration and motion correction of brain images. Neuroimage *17*, 825-841. 10.1016/s1053-8119(02)91132-8.
13. Jenkinson, M., and Smith, S. (2001). A global optimisation method for robust affine registration of brain images. Medical Image Analysis *5*, 143-156. 10.1016/s1361-8415(01)00036-6.

14. Greve, D.N., and Fischl, B. (2009). Accurate and robust brain image alignment using boundary-based registration. Neuroimage *48*, 63-72. 10.1016/j.neuroimage.2009.06.060.
15. Power, J.D., Mitra, A., Laumann, T.O., Snyder, A.Z., Schlaggar, B.L., and Petersen, S.E. (2014). Methods to detect, characterize, and remove motion artifact in resting state fMRI. Neuroimage *84*, 320-341. 10.1016/j.neuroimage.2013.08.048.
16. Behzadi, Y., Restom, K., Liau, J., and Liu, T.T. (2007). A component based noise correction method (CompCor) for BOLD and perfusion based fMRI. Neuroimage *37*, 90-101. 10.1016/j.neuroimage.2007.04.042.
17. Satterthwaite, T.D., Elliott, M.A., Gerraty, R.T., Ruparel, K., Loughead, J., Calkins, M.E., Eickhoff, S.B., Hakonarson, H., Gur, R.C., Gur, R.E., and Wolf, D.H. (2013). An improved framework for confound regression and filtering for control of motion artifact in the preprocessing of resting-state functional connectivity data. Neuroimage *64*, 240-256. 10.1016/j.neuroimage.2012.08.052.
18. Patriat, R., Reynolds, R.C., and Birn, R.M. (2017). An improved model of motion-related signal changes in fMRI. Neuroimage *144*, 74-82. 10.1016/j.neuroimage.2016.08.051.
19. Lanczos, C. (1964). Evaluation of Noisy Data. Journal of the Society for Industrial and Applied Mathematics Series B Numerical Analysis *1*, 76-85. 10.1137/0701007.
20. Abraham, A., Pedregosa, F., Eickenberg, M., Gervais, P., Mueller, A., Kossaifi, J., Gramfort, A., Thirion, B., and Varoquaux, G. (2014). Machine learning for neuroimaging with scikit-learn. Frontiers in neuroinformatics *8*, 14. 10.3389/fninf.2014.00014.
21. Jansma, J.M., Ramsey, N.F., Coppola, R., and Kahn, R.S. (2000). Specific versus nonspecific brain activity in a parametric N-back task. Neuroimage *12*, 688-697. 10.1006/nimg.2000.0645.
22. Glasser, M.F., Coalson, T.S., Robinson, E.C., Hacker, C.D., Harwell, J., Yacoub, E., Ugurbil, K., Andersson, J., Beckmann, C.F., Jenkinson, M., et al. (2016). A multi-modal parcellation of human cerebral cortex. Nature *536*, 171-178. 10.1038/nature18933.
23. Brodt, S., Pöhlchen, D., Flanagin, V.L., Glasauer, S., Gais, S., and Schönauer, M. (2016). Rapid and independent memory formation in the parietal cortex. Proceedings of the National Academy of Sciences *113*, 13251-13256. 10.1073/pnas.1605719113.
24. Critchley, H.D., Mathias, C.J., and Dolan, R.J. (2001). Neural activity in the human brain relating to uncertainty and arousal during anticipation. Neuron *29*, 537-545. 10.1016/s0896-6273(01)00225-2.
25. Knutson, K.M., Wood, J.N., and Grafman, J. (2004). Brain activation in processing temporal sequence: an fMRI study. Neuroimage *23*, 1299-1307. 10.1016/j.neuroimage.2004.08.012.

26. Wagner, A.D., Shannon, B.J., Kahn, I., and Buckner, R.L. (2005). Parietal lobe contributions to episodic memory retrieval. Trends Cogn Sci *9*, 445-453. 10.1016/j.tics.2005.07.001.
27. Cai, M.B., Schuck, N.W., Pillow, J.W., and Niv, Y. (2016). A Bayesian method for reducing bias in neural representational similarity analysis. Proceedings of the 30th International Conference on Neural Information Processing Systems. Curran Associates Inc.
28. Mill, R.D., and Cole, M.W. (2023). Neural representation dynamics reveal computational principles of cognitive task learning. bioRxiv. 10.1101/2023.06.27.546751.
29. Liu, Y., Dolan, R.J., Higgins, C., Penagos, H., Woolrich, M.W., Olafsdottir, H.F., Barry, C., Kurth-Nelson, Z., and Behrens, T.E. (2021). Temporally delayed linear modelling (TDLM) measures replay in both animals and humans. Elife *10*. 10.7554/eLife.66917.
30. Vallesi, A., McIntosh, A.R., Crescentini, C., and Stuss, D.T. (2012). fMRI investigation of speed-accuracy strategy switching. Human brain mapping *33*, 1677-1688. 10.1002/hbm.21312.
31. Kim, C., Cilles, S.E., Johnson, N.F., and Gold, B.T. (2012). Domain general and domain preferential brain regions associated with different types of task switching: a meta-analysis. Human brain mapping *33*, 130-142. 10.1002/hbm.21199.
