## supplementary table S1 for "Cost-benefit Tradeoff Mediates the Rule- to Memory-based Processing Transition during Practice"

**Table S1.1. Rule effect in rule trial**

| HCPex region number | HCPex region name | p(one-tail) | FDR-p(one-tail) |
|---|---|---|---|
| 1 | Primary_Visual_Cortex_L | 0.000188 | 0.002107 |
| 2 | Second_Visual_Area_L | 0.000799 | 0.003573 |
| 3 | Third_Visual_Area_L | 0.000139 | 0.002082 |
| 4 | Fourth_Visual_Area_L | 0.000165 | 0.002107 |
| 5 | IntraParietal_Sulcus_Area_1_L | 0.000244 | 0.002229 |
| 9 | Area_V6A_L | 0.000180 | 0.002107 |
| 11 | Fusiform_Face_Complex_L | 0.000654 | 0.003347 |
| 12 | Posterior_InferoTemporal_complex_L | 0.000217 | 0.002229 |
| 13 | Eighth_Visual_Area_L | 0.000486 | 0.002868 |
| 15 | VentroMedial_Visual_Area_2_L | 0.000050 | 0.001755 |
| 19 | Area_Lateral_Occipital_1_L | 0.000183 | 0.002107 |
| 20 | Area_Lateral_Occipital_2_L | 0.000996 | 0.003620 |
| 23 | Middle_Temporal_Area_L | 0.000073 | 0.001991 |
| 25 | Area_V3CD_L | 0.000985 | 0.003614 |
| 26 | Area_V4t_L | 0.000920 | 0.003587 |
| 31 | Primary_Motor_Cortex_L | 0.000962 | 0.003587 |
| 32 | Area_23c_L | 0.000068 | 0.001991 |
| 37 | Area_5m_ventral_L | 0.000935 | 0.003587 |
| 38 | Area_6m_anterior_L | 0.000893 | 0.003587 |
| 39 | Area_6mp_L | 0.000875 | 0.003587 |
| 40 | Supplementary_and_Cingulate_Eye_Field_L | 0.000259 | 0.002244 |
| 44 | Rostral_Area_6_L | 0.000524 | 0.002914 |
| 45 | Ventral_Area_6_L | 0.000327 | 0.002411 |
| 60 | Auditory_4_Complex_L | 0.000569 | 0.003026 |
| 61 | Auditory_5_Complex_L | 0.000713 | 0.003447 |
| 64 | Area_STSd_posterior_L | 0.000087 | 0.001991 |
| 65 | Area_STSv_anterior_L | 0.000785 | 0.003573 |
| 66 | Area_STSv_posterior_L | 0.000879 | 0.003587 |
| 69 | Anterior_Ventral_Insular_Area_L | 0.000954 | 0.003587 |
| 73 | Area_Frontal_Opercular_5_L | 0.000124 | 0.002016 |
| 75 | Middle_Insular_Area_L | 0.000030 | 0.001640 |
| 76 | Para-Insular_Area_L | 0.000406 | 0.002632 |
| 79 | Posterior_Insular_Area_2_L | 0.000110 | 0.001991 |
| 85 | ParaHippocampal_Area_1_L | 0.000298 | 0.002275 |
| 88 | Area_PHT_L | 0.000239 | 0.002229 |
| 89 | Area_TE1_anterior_L | 0.000139 | 0.002082 |
| 92 | Area_TE2_anterior_L | 0.000087 | 0.001991 |
| 96 | PeriSylvian_Language_Area_L | 0.000589 | 0.003053 |
| 98 | Area_TemporoParietoOccipital_Junction_1_L | 0.000288 | 0.002275 |
| 100 | Area_TemporoParietoOccipital_Junction_3_L | 0.000929 | 0.003587 |
| 102 | Medial_Area_7A_L | 0.000195 | 0.002107 |
| 104 | Lateral_Area_7P_L | 0.000812 | 0.003573 |
| 105 | Medial_Area_7P_L | 0.000509 | 0.002914 |
| 107 | Area_Lateral_IntraParietal_dorsal_L | 0.000113 | 0.001991 |

| | | | |
|---|---|---|---|
| 108 | Area_Lateral_IntraParietal_ventral_L | 0.000194 | 0.002107 |
| 109 | Medial_IntraParietal_Area_L | 0.000962 | 0.003587 |
| 110 | Ventral_IntraParietal_Complex_L | 0.000104 | 0.001991 |
| 111 | Area_IntraParietal_0_L | 0.000035 | 0.001640 |
| 113 | Area_IntraParietal_2_L | 0.000894 | 0.003587 |
| 119 | Area_PGp_L | 0.000470 | 0.002868 |
| 127 | Dorsal_Transitional_Visual_Area_L | 0.000391 | 0.002604 |
| 128 | PreCuneus_Visual_Area_L | 0.000028 | 0.001640 |
| 130 | Parieto-Occipital_Sulcus_Area_2_L | 0.000576 | 0.003026 |
| 132 | RetroSplenial_Complex_L | 0.000753 | 0.003489 |
| 138 | Area_8BM_L | 0.000172 | 0.002107 |
| 142 | Area_anterior_32_prime_L | 0.000042 | 0.001640 |
| 143 | Area_dorsal_32_L | 0.000335 | 0.002411 |
| 144 | Area_posterior_24_L | 0.000520 | 0.002914 |
| 147 | Area_p32_prime_L | 0.000507 | 0.002914 |
| 159 | Area_44_L | 0.000293 | 0.002275 |
| 162 | Area_anterior_47r_L | 0.000675 | 0.003411 |
| 165 | Area_IFSa_L | 0.000023 | 0.001640 |
| 166 | Area_IFSp_L | 0.000283 | 0.002275 |
| 168 | Area_46_L | 0.000429 | 0.002738 |
| 172 | Area_8C_L | 0.000265 | 0.002244 |
| 173 | Area_9-46d_L | 0.000118 | 0.001991 |
| 175 | Area_9_Posterior_L | 0.000002 | 0.000971 |
| 176 | Area_anterior_9-46v_L | 0.000233 | 0.002229 |
| 181 | Primary_Visual_Cortex_R | 0.000092 | 0.001991 |
| 184 | Fourth_Visual_Area_R | 0.000395 | 0.002604 |
| 191 | Fusiform_Face_Complex_R | 0.000274 | 0.002269 |
| 195 | VentroMedial_Visual_Area_2_R | 0.000009 | 0.001212 |
| 197 | Ventral_Visual_Complex_R | 0.000333 | 0.002411 |
| 205 | Area_V3CD_R | 0.000472 | 0.002868 |
| 213 | Dorsal_Area_24d_R | 0.000487 | 0.002868 |
| 219 | Area_6mp_R | 0.000375 | 0.002604 |
| 222 | Area_6_anterior_R | 0.000246 | 0.002229 |
| 224 | Rostral_Area_6_R | 0.000704 | 0.003447 |
| 227 | Premotor_Eye_Field_R | 0.000239 | 0.002229 |
| 249 | Anterior_Ventral_Insular_Area_R | 0.000175 | 0.002107 |
| 252 | Frontal_Opercular_Area_4_R | 0.000170 | 0.002107 |
| 253 | Area_Frontal_Opercular_5_R | 0.000939 | 0.003587 |
| 264 | Area_TF_R | 0.000026 | 0.001640 |
| 267 | ParaHippocampal_Area_3_R | 0.000236 | 0.002229 |
| 273 | Area_TE2_posterior_R | 0.000118 | 0.001991 |
| 281 | Lateral_Area_7A_R | 0.000840 | 0.003587 |
| 288 | Area_Lateral_IntraParietal_ventral_R | 0.000097 | 0.001991 |
| 290 | Ventral_IntraParietal_Complex_R | 0.000726 | 0.003447 |
| 293 | Area_IntraParietal_2_R | 0.000968 | 0.003587 |
| 299 | Area_PGp_R | 0.000388 | 0.002604 |
| 301 | Area_23d_R | 0.000573 | 0.003026 |

| | | | |
|---|---|---|---|
| 309 | Parieto-Occipital_Sulcus_Area_1_R | 0.000932 | 0.003587 |
| 310 | Parieto-Occipital_Sulcus_Area_2_R | 0.000539 | 0.002952 |
| 318 | Area_8BM_R | 0.000058 | 0.001893 |
| 319 | Area_9_Middle_R | 0.000727 | 0.003447 |
| 326 | Area_p32_R | 0.000861 | 0.003587 |
| 335 | Area_47s_R | 0.000752 | 0.003489 |
| 339 | Area_44_R | 0.000817 | 0.003573 |
| 340 | Area_45_R | 0.000150 | 0.002107 |
| 341 | Area_47l_(47_lateral)_R | 0.000263 | 0.002244 |
| 344 | Area_IFJp_R | 0.000355 | 0.002511 |
| 345 | Area_IFSa_R | 0.000442 | 0.002773 |
| 349 | Area_8Ad_R | 0.000944 | 0.003587 |
| 350 | Area_8Av_R | 0.000691 | 0.003445 |
| 353 | Area_9-46d_R | 0.000009 | 0.001212 |
| 382 | Putamen_L | 0.000041 | 0.001640 |
| 415 | Putamen_R | 0.000817 | 0.003573 |

**Table S1.2. Rule effect in memory trial**

| HCPex region number | HCPex region name | p(one-tail) | FDR-p(one-tail) |
|---|---|---|---|
| Null | Null | Null | Null |

**Table S1.3. Higher Rule effect in rule than memory trials**

| HCPex region number | HCPex region name | p(one-tail) | FDR-p(one-tail) |
|---|---|---|---|
| regionNo | Name | p | q |
| 1 | Primary_Visual_Cortex_L | 0.000170 | 0.001336 |
| 2 | Second_Visual_Area_L | 0.000249 | 0.001686 |
| 3 | Third_Visual_Area_L | 0.000015 | 0.000456 |
| 4 | Fourth_Visual_Area_L | 0.000015 | 0.000456 |
| 5 | IntraParietal_Sulcus_Area_1_L | 0.000046 | 0.000742 |
| 9 | Area_V6A_L | 0.000516 | 0.002257 |
| 11 | Fusiform_Face_Complex_L | 0.000034 | 0.000652 |
| 13 | Eighth_Visual_Area_L | 0.000281 | 0.001711 |
| 15 | VentroMedial_Visual_Area_2_L | 0.000004 | 0.000394 |
| 17 | Ventral_Visual_Complex_L | 0.000481 | 0.002257 |
| 20 | Area_Lateral_Occipital_2_L | 0.000845 | 0.003005 |
| 24 | Area_PH_L | 0.000437 | 0.002126 |
| 26 | Area_V4t_L | 0.000567 | 0.002424 |
| 28 | Area_2_L | 0.000412 | 0.002080 |
| 30 | Primary_Sensory_Cortex_L | 0.000715 | 0.002764 |
| 31 | Primary_Motor_Cortex_L | 0.000021 | 0.000519 |
| 33 | Dorsal_Area_24d_L | 0.000761 | 0.002875 |
| 35 | Area_5L_L | 0.000175 | 0.001336 |
| 40 | Supplementary_and_Cingulate_Eye_Field_L | 0.000174 | 0.001336 |
| 42 | Area_6_anterior_L | 0.000417 | 0.002080 |
| 43 | Dorsal_area_6_L | 0.000014 | 0.000456 |
| 44 | Rostral_Area_6_L | 0.000355 | 0.001960 |
| 45 | Ventral_Area_6_L | 0.000011 | 0.000456 |

| | | | |
|---|---|---|---|
| 46 | Frontal_Eye_Fields_L | 0.000011 | 0.000456 |
| 56 | Medial_Belt_Complex_L | 0.000122 | 0.001076 |
| 57 | ParaBelt_Complex_L | 0.000048 | 0.000742 |
| 60 | Auditory_4_Complex_L | 0.000118 | 0.001066 |
| 61 | Auditory_5_Complex_L | 0.000033 | 0.000652 |
| 64 | Area_STSd_posterior_L | 0.000633 | 0.002560 |
| 66 | Area_STSv_posterior_L | 0.000977 | 0.003248 |
| 67 | Area_TA2_L | 0.000117 | 0.001066 |
| 68 | Anterior_Agranular_Insula_Complex_L | 0.000988 | 0.003258 |
| 72 | Frontal_Opercular_Area_4_L | 0.000402 | 0.002080 |
| 75 | Middle_Insular_Area_L | 0.000512 | 0.002257 |
| 79 | Posterior_Insular_Area_2_L | 0.000003 | 0.000352 |
| 80 | Hippocampus_L | 0.000818 | 0.002973 |
| 81 | PreSubiculum_L | 0.000187 | 0.001396 |
| 88 | Area_PHT_L | 0.000068 | 0.000825 |
| 89 | Area_TE1_anterior_L | 0.000012 | 0.000456 |
| 90 | Area_TE1_Middle_L | 0.000273 | 0.001686 |
| 92 | Area_TE2_anterior_L | 0.000052 | 0.000742 |
| 93 | Area_TE2_posterior_L | 0.000768 | 0.002875 |
| 94 | Area_TG_dorsal_L | 0.000618 | 0.002560 |
| 97 | Superior_Temporal_Visual_Area_L | 0.000064 | 0.000805 |
| 98 | Area_TemporoParietoOccipital_Junction_1_L | 0.000064 | 0.000805 |
| 102 | Medial_Area_7A_L | 0.000079 | 0.000879 |
| 104 | Lateral_Area_7P_L | 0.000506 | 0.002257 |
| 105 | Medial_Area_7P_L | 0.000833 | 0.003001 |
| 106 | Anterior_IntraParietal_Area_L | 0.000638 | 0.002560 |
| 107 | Area_Lateral_IntraParietal_dorsal_L | 0.000002 | 0.000352 |
| 108 | Area_Lateral_IntraParietal_ventral_L | 0.000070 | 0.000825 |
| 109 | Medial_IntraParietal_Area_L | 0.000097 | 0.000970 |
| 110 | Ventral_IntraParietal_Complex_L | 0.000015 | 0.000456 |
| 111 | Area_IntraParietal_0_L | 0.000038 | 0.000702 |
| 112 | Area_IntraParietal_1_L | 0.000504 | 0.002257 |
| 113 | Area_IntraParietal_2_L | 0.000273 | 0.001686 |
| 114 | Area_PF_Complex_L | 0.000535 | 0.002315 |
| 118 | Area_PGi_L | 0.000270 | 0.001686 |
| 119 | Area_PGp_L | 0.000104 | 0.001011 |
| 120 | Area_PGs_L | 0.000891 | 0.003067 |
| 123 | Area_31pd_L | 0.000270 | 0.001686 |
| 128 | PreCuneus_Visual_Area_L | 0.000014 | 0.000456 |
| 129 | Parieto-Occipital_Sulcus_Area_1_L | 0.000936 | 0.003156 |
| 130 | Parieto-Occipital_Sulcus_Area_2_L | 0.000018 | 0.000456 |
| 132 | RetroSplenial_Complex_L | 0.000091 | 0.000968 |
| 138 | Area_8BM_L | 0.000294 | 0.001756 |
| 142 | Area_anterior_32_prime_L | 0.000003 | 0.000352 |
| 143 | Area_dorsal_32_L | 0.000869 | 0.003041 |
| 147 | Area_p32_prime_L | 0.000314 | 0.001826 |
| 159 | Area_44_L | 0.000623 | 0.002560 |

| | | | |
|---|---|---|---|
| 160 | Area_45_L | 0.000487 | 0.002257 |
| 165 | Area_IFSa_L | 0.000094 | 0.000968 |
| 166 | Area_IFSp_L | 0.000391 | 0.002074 |
| 168 | Area_46_L | 0.000395 | 0.002074 |
| 172 | Area_8C_L | 0.000017 | 0.000456 |
| 173 | Area_9-46d_L | 0.000899 | 0.003067 |
| 175 | Area_9_Posterior_L | 0.000095 | 0.000968 |
| 177 | Inferior_6-8_Transitional_Area_L | 0.000192 | 0.001412 |
| 178 | Area_posterior_9-46v_L | 0.000131 | 0.001104 |
| 181 | Primary_Visual_Cortex_R | 0.000010 | 0.000456 |
| 183 | Third_Visual_Area_R | 0.000718 | 0.002764 |
| 184 | Fourth_Visual_Area_R | 0.000321 | 0.001836 |
| 185 | IntraParietal_Sulcus_Area_1_R | 0.000769 | 0.002875 |
| 187 | Area_V3B_R | 0.000706 | 0.002764 |
| 191 | Fusiform_Face_Complex_R | 0.000057 | 0.000768 |
| 195 | VentroMedial_Visual_Area_2_R | 0.000053 | 0.000742 |
| 197 | Ventral_Visual_Complex_R | 0.000633 | 0.002560 |
| 199 | Area_Lateral_Occipital_1_R | 0.000473 | 0.002257 |
| 205 | Area_V3CD_R | 0.000348 | 0.001960 |
| 221 | Area_55b_R | 0.000427 | 0.002101 |
| 222 | Area_6_anterior_R | 0.000161 | 0.001333 |
| 226 | Frontal_Eye_Fields_R | 0.000298 | 0.001756 |
| 227 | Premotor_Eye_Field_R | 0.000247 | 0.001686 |
| 235 | Lateral_Belt_Complex_R | 0.000025 | 0.000565 |
| 236 | Medial_Belt_Complex_R | 0.000223 | 0.001576 |
| 252 | Frontal_Opercular_Area_4_R | 0.000358 | 0.001960 |
| 255 | Middle_Insular_Area_R | 0.000941 | 0.003156 |
| 257 | Pirform_Cortex_R | 0.000596 | 0.002521 |
| 264 | Area_TF_R | 0.000507 | 0.002257 |
| 273 | Area_TE2_posterior_R | 0.000053 | 0.000742 |
| 278 | Area_TemporoParietoOccipital_Junction_1_R | 0.000501 | 0.002257 |
| 288 | Area_Lateral_IntraParietal_ventral_R | 0.000026 | 0.000565 |
| 305 | Area_7m_R | 0.000795 | 0.002917 |
| 309 | Parieto-Occipital_Sulcus_Area_1_R | 0.000387 | 0.002074 |
| 310 | Parieto-Occipital_Sulcus_Area_2_R | 0.000126 | 0.001090 |
| 312 | RetroSplenial_Complex_R | 0.000788 | 0.002917 |
| 319 | Area_9_Middle_R | 0.000875 | 0.003041 |
| 322 | Area_anterior_32_prime_R | 0.000850 | 0.003005 |
| 335 | Area_47s_R | 0.000268 | 0.001686 |
| 345 | Area_IFSa_R | 0.000417 | 0.002080 |
| 350 | Area_8Av_R | 0.000040 | 0.000715 |
| 353 | Area_9-46d_R | 0.000209 | 0.001502 |
| 357 | Inferior_6-8_Transitional_Area_R | 0.000253 | 0.001686 |
| 358 | Area_posterior_9-46v_R | 0.000116 | 0.001066 |
| 382 | Putamen_L | 0.000052 | 0.000742 |
| 383 | Caudate_L | 0.000676 | 0.002685 |
| 415 | Putamen_R | 0.000073 | 0.000832 |

| | | | |
|---|---|---|---|
| 416 | Caudate_R | 0.000175 | 0.001336 |

**Table S1.4. Cue effect in rule trial**

| HCPex region number | HCPex region name | p(one-tail) | FDR-p(one-tail) |
|---|---|---|---|
| 27 | Area_1_L | 0.000005 | 0.000672 |
| 30 | Primary_Sensory_Cortex_L | 0.000441 | 0.028613 |
| 31 | Primary_Motor_Cortex_L | 0.000674 | 0.037410 |
| 43 | Dorsal_area_6_L | 0.000769 | 0.037410 |
| 207 | Area_1_R | 0.000005 | 0.000672 |
| 208 | Area_2_R | 0.000123 | 0.009582 |
| 209 | Area_3a_R | 0.000042 | 0.004105 |
| 210 | Primary_Sensory_Cortex_R | 0.000002 | 0.000672 |

**Table S1.5. Cue effect in memory trial**

| HCPex region number | HCPex region name | p(one-tail) | FDR-p(one-tail) |
|---|---|---|---|
| 1 | Primary_Visual_Cortex_L | 0.000000 | 0.000012 |
| 2 | Second_Visual_Area_L | 0.000000 | 0.000005 |
| 3 | Third_Visual_Area_L | 0.000003 | 0.000048 |
| 4 | Fourth_Visual_Area_L | 0.000002 | 0.000031 |
| 11 | Fusiform_Face_Complex_L | 0.000074 | 0.000494 |
| 17 | Ventral_Visual_Complex_L | 0.000454 | 0.001900 |
| 24 | Area_PH_L | 0.000782 | 0.002714 |
| 27 | Area_1_L | 0.000001 | 0.000023 |
| 28 | Area_2_L | 0.000014 | 0.000166 |
| 29 | Area_3a_L | 0.000033 | 0.000295 |
| 30 | Primary_Sensory_Cortex_L | 0.000000 | 0.000002 |
| 31 | Primary_Motor_Cortex_L | 0.000000 | 0.000001 |
| 33 | Dorsal_Area_24d_L | 0.000001 | 0.000016 |
| 35 | Area_5L_L | 0.000141 | 0.000794 |
| 38 | Area_6m_anterior_L | 0.000960 | 0.003166 |
| 40 | Supplementary_and_Cingulate_Eye_Field_L | 0.000025 | 0.000233 |
| 41 | Area_55b_L | 0.000170 | 0.000934 |
| 42 | Area_6_anterior_L | 0.000136 | 0.000791 |
| 43 | Dorsal_area_6_L | 0.000017 | 0.000186 |
| 44 | Rostral_Area_6_L | 0.000906 | 0.003037 |
| 45 | Ventral_Area_6_L | 0.000060 | 0.000432 |
| 46 | Frontal_Eye_Fields_L | 0.000002 | 0.000037 |
| 48 | Area_43_L | 0.000041 | 0.000352 |
| 50 | Area_OP1-SII_L | 0.000051 | 0.000381 |
| 51 | Area_OP2-3-VS_L | 0.000633 | 0.002436 |
| 52 | Area_OP4-PV_L | 0.000613 | 0.002386 |
| 74 | Insular_Granular_Complex_L | 0.000198 | 0.001058 |
| 88 | Area_PHT_L | 0.000048 | 0.000381 |
| 93 | Area_TE2_posterior_L | 0.000447 | 0.001891 |
| 98 | Area_TemporoParietoOccipital_Junction_1_L | 0.000295 | 0.001382 |
| 102 | Medial_Area_7A_L | 0.000133 | 0.000784 |
| 103 | Area_7PC_L | 0.000017 | 0.000186 |

| | | | |
|---|---|---|---|
| 106 | Anterior_IntraParietal_Area_L | 0.000085 | 0.000551 |
| 107 | Area_Lateral_IntraParietal_dorsal_L | 0.000392 | 0.001694 |
| 108 | Area_Lateral_IntraParietal_ventral_L | 0.000002 | 0.000031 |
| 110 | Ventral_IntraParietal_Complex_L | 0.000603 | 0.002371 |
| 111 | Area_IntraParietal_0_L | 0.000559 | 0.002265 |
| 112 | Area_IntraParietal_1_L | 0.000654 | 0.002447 |
| 113 | Area_IntraParietal_2_L | 0.000286 | 0.001355 |
| 114 | Area_PF_Complex_L | 0.000000 | 0.000000 |
| 115 | Area_PFm_Complex_L | 0.000001 | 0.000023 |
| 116 | Area_PF_Opercular_L | 0.000332 | 0.001501 |
| 117 | Area_PFt_L | 0.000225 | 0.001166 |
| 119 | Area_PGp_L | 0.000005 | 0.000070 |
| 120 | Area_PGs_L | 0.000279 | 0.001355 |
| 121 | Area_23d_L | 0.000678 | 0.002512 |
| 122 | Area_31a_L | 0.000175 | 0.000945 |
| 123 | Area_31pd_L | 0.000060 | 0.000432 |
| 124 | Area_31p_ventral_L | 0.000130 | 0.000778 |
| 125 | Area_7m_L | 0.000020 | 0.000200 |
| 126 | Area_dorsal_23_a+b_L | 0.000697 | 0.002535 |
| 128 | PreCuneus_Visual_Area_L | 0.000046 | 0.000381 |
| 130 | Parieto-Occipital_Sulcus_Area_2_L | 0.000003 | 0.000052 |
| 142 | Area_anterior_32_prime_L | 0.000652 | 0.002447 |
| 159 | Area_44_L | 0.000789 | 0.002715 |
| 160 | Area_45_L | 0.000020 | 0.000200 |
| 161 | Area_47l_(47_lateral)_L | 0.000874 | 0.002982 |
| 165 | Area_IFSa_L | 0.000116 | 0.000706 |
| 172 | Area_8C_L | 0.000215 | 0.001132 |
| 177 | Inferior_6-8_Transitional_Area_L | 0.000088 | 0.000552 |
| 178 | Area_posterior_9-46v_L | 0.000504 | 0.002063 |
| 181 | Primary_Visual_Cortex_R | 0.000000 | 0.000000 |
| 182 | Second_Visual_Area_R | 0.000000 | 0.000002 |
| 183 | Third_Visual_Area_R | 0.000002 | 0.000037 |
| 184 | Fourth_Visual_Area_R | 0.000002 | 0.000032 |
| 185 | IntraParietal_Sulcus_Area_1_R | 0.000367 | 0.001621 |
| 188 | Sixth_Visual_Area_R | 0.000005 | 0.000064 |
| 190 | Seventh_Visual_Area_R | 0.000426 | 0.001823 |
| 192 | Posterior_InferoTemporal_complex_R | 0.000731 | 0.002588 |
| 193 | Eighth_Visual_Area_R | 0.000049 | 0.000381 |
| 196 | VentroMedial_Visual_Area_3_R | 0.000007 | 0.000090 |
| 199 | Area_Lateral_Occipital_1_R | 0.000078 | 0.000517 |
| 200 | Area_Lateral_Occipital_2_R | 0.000599 | 0.002371 |
| 203 | Middle_Temporal_Area_R | 0.000994 | 0.003230 |
| 205 | Area_V3CD_R | 0.000235 | 0.001203 |
| 206 | Area_V4t_R | 0.000285 | 0.001355 |
| 207 | Area_1_R | 0.000000 | 0.000013 |
| 208 | Area_2_R | 0.000004 | 0.000062 |
| 209 | Area_3a_R | 0.000000 | 0.000004 |

| | | | |
|---|---|---|---|
| 210 | Primary_Sensory_Cortex_R | 0.000000 | 0.000003 |
| 211 | Primary_Motor_Cortex_R | 0.000001 | 0.000015 |
| 213 | Dorsal_Area_24d_R | 0.000000 | 0.000013 |
| 215 | Area_5L_R | 0.000247 | 0.001248 |
| 216 | Area_5m_R | 0.000707 | 0.002548 |
| 218 | Area_6m_anterior_R | 0.000376 | 0.001645 |
| 219 | Area_6mp_R | 0.000068 | 0.000472 |
| 223 | Dorsal_area_6_R | 0.000018 | 0.000186 |
| 225 | Ventral_Area_6_R | 0.000107 | 0.000658 |
| 226 | Frontal_Eye_Fields_R | 0.000001 | 0.000015 |
| 230 | Area_OP1-SII_R | 0.000000 | 0.000000 |
| 258 | Area_Posterior_Insular_1_R | 0.000732 | 0.002588 |
| 266 | ParaHippocampal_Area_2_R | 0.000256 | 0.001275 |
| 278 | Area_TemporoParietoOccipital_Junction_1_R | 0.000015 | 0.000182 |
| 280 | Area_TemporoParietoOccipital_Junction_3_R | 0.000156 | 0.000868 |
| 281 | Lateral_Area_7A_R | 0.000648 | 0.002447 |
| 282 | Medial_Area_7A_R | 0.000341 | 0.001523 |
| 283 | Area_7PC_R | 0.000044 | 0.000372 |
| 284 | Lateral_Area_7P_R | 0.000026 | 0.000237 |
| 286 | Anterior_IntraParietal_Area_R | 0.000687 | 0.002522 |
| 287 | Area_Lateral_IntraParietal_dorsal_R | 0.000467 | 0.001933 |
| 288 | Area_Lateral_IntraParietal_ventral_R | 0.000061 | 0.000432 |
| 289 | Medial_IntraParietal_Area_R | 0.000000 | 0.000000 |
| 291 | Area_IntraParietal_0_R | 0.000276 | 0.001355 |
| 292 | Area_IntraParietal_1_R | 0.000317 | 0.001449 |
| 293 | Area_IntraParietal_2_R | 0.000004 | 0.000062 |
| 294 | Area_PF_Complex_R | 0.000051 | 0.000381 |
| 295 | Area_PFm_Complex_R | 0.000029 | 0.000266 |
| 300 | Area_PGs_R | 0.000017 | 0.000186 |
| 302 | Area_31a_R | 0.000739 | 0.002589 |
| 304 | Area_31p_ventral_R | 0.000310 | 0.001435 |
| 305 | Area_7m_R | 0.000595 | 0.002371 |
| 306 | Area_dorsal_23_a+b_R | 0.000012 | 0.000148 |
| 310 | Parieto-Occipital_Sulcus_Area_2_R | 0.000893 | 0.003022 |
| 348 | Area_46_R | 0.000921 | 0.003061 |
| 350 | Area_8Av_R | 0.000021 | 0.000200 |
| 357 | Inferior_6-8_Transitional_Area_R | 0.000086 | 0.000551 |
| 358 | Area_posterior_9-46v_R | 0.000050 | 0.000381 |
| 381 | Thal_Ventral_posterolateral_L | 0.000139 | 0.000794 |
| 382 | Putamen_L | 0.000074 | 0.000494 |
| 415 | Putamen_R | 0.000998 | 0.003230 |

**Table S1.6. Higher Cue effect in memory than rule trials**

| HCPex region number | HCPex region name | p(one-tail) | FDR-p(one-tail) |
|---|---|---|---|
| 2 | Second_Visual_Area_L | 0.000405 | 0.003941 |
| 3 | Third_Visual_Area_L | 0.000080 | 0.001475 |
| 4 | Fourth_Visual_Area_L | 0.000001 | 0.000110 |

| | | | |
|---|---|---|---|
| 5 | IntraParietal_Sulcus_Area_1_L | 0.000903 | 0.006388 |
| 11 | Fusiform_Face_Complex_L | 0.000084 | 0.001477 |
| 15 | VentroMedial_Visual_Area_2_L | 0.000019 | 0.000695 |
| 26 | Area_V4t_L | 0.000764 | 0.005943 |
| 35 | Area_5L_L | 0.000201 | 0.002796 |
| 44 | Rostral_Area_6_L | 0.000661 | 0.005382 |
| 49 | Frontal_Opercular_Area_1_L | 0.000274 | 0.003553 |
| 64 | Area_STSd_posterior_L | 0.000033 | 0.000906 |
| 81 | PreSubiculum_L | 0.000080 | 0.001475 |
| 85 | ParaHippocampal_Area_1_L | 0.000181 | 0.002604 |
| 88 | Area_PHT_L | 0.000119 | 0.002013 |
| 93 | Area_TE2_posterior_L | 0.000664 | 0.005382 |
| 98 | Area_TemporoParietoOccipital_Junction_1_L | 0.000173 | 0.002581 |
| 103 | Area_7PC_L | 0.000222 | 0.002974 |
| 107 | Area_Lateral_IntraParietal_dorsal_L | 0.000702 | 0.005573 |
| 108 | Area_Lateral_IntraParietal_ventral_L | 0.000000 | 0.000110 |
| 110 | Ventral_IntraParietal_Complex_L | 0.000003 | 0.000302 |
| 111 | Area_IntraParietal_0_L | 0.000445 | 0.004123 |
| 119 | Area_PGp_L | 0.000326 | 0.003941 |
| 122 | Area_31a_L | 0.000171 | 0.002581 |
| 125 | Area_7m_L | 0.000658 | 0.005382 |
| 128 | PreCuneus_Visual_Area_L | 0.000006 | 0.000466 |
| 130 | Parieto-Occipital_Sulcus_Area_2_L | 0.000885 | 0.006388 |
| 142 | Area_anterior_32_prime_L | 0.000436 | 0.004123 |
| 160 | Area_45_L | 0.000015 | 0.000695 |
| 165 | Area_IFSa_L | 0.000031 | 0.000906 |
| 166 | Area_IFSp_L | 0.000003 | 0.000302 |
| 172 | Area_8C_L | 0.000901 | 0.006388 |
| 178 | Area_posterior_9-46v_L | 0.000377 | 0.003941 |
| 181 | Primary_Visual_Cortex_R | 0.000015 | 0.000695 |
| 182 | Second_Visual_Area_R | 0.000021 | 0.000695 |
| 184 | Fourth_Visual_Area_R | 0.000050 | 0.001206 |
| 185 | IntraParietal_Sulcus_Area_1_R | 0.000315 | 0.003941 |
| 191 | Fusiform_Face_Complex_R | 0.000393 | 0.003941 |
| 193 | Eighth_Visual_Area_R | 0.000378 | 0.003941 |
| 195 | VentroMedial_Visual_Area_2_R | 0.000021 | 0.000695 |
| 196 | VentroMedial_Visual_Area_3_R | 0.000522 | 0.004509 |
| 197 | Ventral_Visual_Complex_R | 0.000017 | 0.000695 |
| 199 | Area_Lateral_Occipital_1_R | 0.000055 | 0.001267 |
| 200 | Area_Lateral_Occipital_2_R | 0.000501 | 0.004434 |
| 202 | Medial_Superior_Temporal_Area_R | 0.000992 | 0.006727 |
| 204 | Area_PH_R | 0.000353 | 0.003941 |
| 227 | Premotor_Eye_Field_R | 0.000362 | 0.003941 |
| 282 | Medial_Area_7A_R | 0.000369 | 0.003941 |
| 284 | Lateral_Area_7P_R | 0.000156 | 0.002529 |
| 288 | Area_Lateral_IntraParietal_ventral_R | 0.000043 | 0.001126 |
| 289 | Medial_IntraParietal_Area_R | 0.000020 | 0.000695 |

| | | | |
|---|---|---|---|
| 290 | Ventral_IntraParietal_Complex_R | 0.000463 | 0.004188 |
| 291 | Area_IntraParietal_0_R | 0.000073 | 0.001475 |
| 300 | Area_PGs_R | 0.000841 | 0.006388 |
| 312 | RetroSplenial_Complex_R | 0.000900 | 0.006388 |
| 379 | Thal_Ventral_Lateral_Anterior_L | 0.000400 | 0.003941 |
| 382 | Putamen_L | 0.000078 | 0.001475 |

**Table S1.7. Double dissociation effect (Conjunction)**

| HCPex region number | HCPex region name | p(one-tail) | FDR-p(one-tail) |
|---|---|---|---|
| 2 | Second_Visual_Area_L | 0.000010 | 0.000085 |
| 3 | Third_Visual_Area_L | 0.000000 | 0.000011 |
| 4 | Fourth_Visual_Area_L | 0.000000 | 0.000008 |
| 5 | IntraParietal_Sulcus_Area_1_L | 0.000003 | 0.000040 |
| 11 | Fusiform_Face_Complex_L | 0.000001 | 0.000019 |
| 15 | VentroMedial_Visual_Area_2_L | 0.000000 | 0.000006 |
| 26 | Area_V4t_L | 0.000037 | 0.000194 |
| 35 | Area_5L_L | 0.000006 | 0.000059 |
| 44 | Rostral_Area_6_L | 0.000018 | 0.000125 |
| 64 | Area_STSd_posterior_L | 0.000004 | 0.000044 |
| 81 | PreSubiculum_L | 0.000003 | 0.000043 |
| 88 | Area_PHT_L | 0.000002 | 0.000026 |
| 93 | Area_TE2_posterior_L | 0.000045 | 0.000223 |
| 98 | Area_TemporoParietoOccipital_Junction_1_L | 0.000001 | 0.000018 |
| 107 | Area_Lateral_IntraParietal_dorsal_L | 0.000000 | 0.000008 |
| 108 | Area_Lateral_IntraParietal_ventral_L | 0.000001 | 0.000016 |
| 110 | Ventral_IntraParietal_Complex_L | 0.000000 | 0.000008 |
| 111 | Area_IntraParietal_0_L | 0.000001 | 0.000023 |
| 119 | Area_PGp_L | 0.000004 | 0.000044 |
| 128 | PreCuneus_Visual_Area_L | 0.000000 | 0.000008 |
| 130 | Parieto-Occipital_Sulcus_Area_2_L | 0.000001 | 0.000016 |
| 142 | Area_anterior_32_prime_L | 0.000000 | 0.000007 |
| 160 | Area_45_L | 0.000009 | 0.000081 |
| 165 | Area_IFSa_L | 0.000001 | 0.000016 |
| 166 | Area_IFSp_L | 0.000003 | 0.000041 |
| 172 | Area_8C_L | 0.000001 | 0.000016 |
| 178 | Area_posterior_9-46v_L | 0.000004 | 0.000048 |
| 181 | Primary_Visual_Cortex_R | 0.000000 | 0.000006 |
| 184 | Fourth_Visual_Area_R | 0.000010 | 0.000085 |
| 185 | IntraParietal_Sulcus_Area_1_R | 0.000030 | 0.000172 |
| 191 | Fusiform_Face_Complex_R | 0.000002 | 0.000036 |
| 195 | VentroMedial_Visual_Area_2_R | 0.000001 | 0.000016 |
| 197 | Ventral_Visual_Complex_R | 0.000014 | 0.000108 |
| 199 | Area_Lateral_Occipital_1_R | 0.000017 | 0.000122 |
| 227 | Premotor_Eye_Field_R | 0.000014 | 0.000109 |
| 288 | Area_Lateral_IntraParietal_ventral_R | 0.000000 | 0.000013 |
| 312 | RetroSplenial_Complex_R | 0.000042 | 0.000213 |
| 382 | Putamen_L | 0.000001 | 0.000018 |
